# Combining metabolomics and experimental evolution reveals key mechanisms underlying longevity differences in laboratory evolved *Drosophila melanogaster* populations

**DOI:** 10.1101/2021.10.16.464668

**Authors:** Mark A. Phillips, Kenneth R. Arnold, Zer Vue, Heather K. Beasley, Edgar Garza Lopez, Andrea G. Marshall, Derrick J. Morton, Melanie R. McReynolds, Thomas T. Barter, Antentor Hinton

## Abstract

Experimental evolution with *Drosophila melanogaster* has been used extensively for decades to study aging and longevity. In recent years, the addition of DNA and RNA sequencing to this framework has allowed researchers to leverage the statistical power inherent to experimental evolution study the genetic basis of longevity itself. Here we incorporated metabolomic data into to this framework to generate even deeper insights into the physiological and genetic mechanisms underlying longevity differences in three groups of experimentally evolved *D. melanogaster* populations with different aging and longevity patterns. Our metabolomic analysis found that aging alters mitochondrial metabolism through increased consumption of NAD^+^ and increased usage of the TCA cycle. Combining our genomic and metabolomic data produced a list of biologically relevant candidate genes. Among these candidates, we found significant enrichment for genes and pathways associated with neurological development and function, and carbohydrate metabolism. While we do not explicitly find enrichment for aging canonical genes, neurological dysregulation and carbohydrate metabolism are both known to be associated with accelerated aging and reduced longevity. Taken together, our results provide plausible genetic mechanisms for what might be driving longevity differences in this experimental system. More broadly, our findings demonstrate the value of combining multiple types of omic data with experimental evolution when attempting to dissect mechanisms underlying complex and highly polygenic traits like aging.

## Introduction

Understanding the factors that drive differences in life history between individuals, populations, and species is a major area of interest for evolutionary biologists ^1–4^. Within this area of research, experimental evolution is a powerful tool for addressing fundamental questions about how natural selection shapes life history. *Drosophila melanogaster* is an extensively used, powerful, genetically tractable model system for studying life histories. For instance, there is a large body of work devoted to subjecting *D. melanogaster* populations to selection regimes that target variables thought to be important in life history evolution to test theoretical predictions ^5–9^. Comparisons between populations selected for radically different life histories have also been successfully used to study the physiological and genetic mechanisms underlying observed differences in traits like longevity, developmental rates, and reproductive output ^10^. Here we are primarily interested in longevity.

Many studies using experimental evolution to study physiological mechanisms underlying longevity differences have focused on the relationship between stress tolerance and life span. For instance, selection for decreased rates of senescence and increased longevity have repeatably been shown to associate with improved desiccation and starvation resistance and vice-versa ^11–15^. Physiological assays have then shown that differences in longevity associated stress tolerances in these systems are tied to differences in lipid, glycogen, and water content ^15–19^. However, deeper, more fine-scale investigations into what other physiological factors are driving differences in these types of experimental systems are currently lacking.

More recently, combining experimental evolution with next-generation sequencing technologies, termed “evolve and resequence” or “E&R”, has emerged as a powerful general tool for dissecting the underlying genetic architecture of complex traits ^20,21^. Using this framework, evolutionary biologists can study the genetic basis of longevity by simply comparing patterns of genetic differentiation between groups of evolved populations with significant differences in mean longevity. Through these comparisons, statistical associations can be made between genetic variants and longevity or related phenotypes. Here it is important to note that using evolution to drive phenotypes in opposite directions generates increased statistical power when making these associations ^22^. Combined with the population-level replication featured in most studies, this approach allows for more powerful statistical inferences to be drawn than what is typically seen in conventional genome-wide association studies ^23^.

At present, several genomic ^15,24–27^ and transcriptomic ^28,29^ E&R studies have generated insights into the genetics of increased longevity and related stress resistances. The integration of other types of “omic” data into this framework can potentially yield similarly powerful insights. Here, we aim to do so by incorporating metabolomic data into the E&R framework. Specifically, we explore metabolomic differences between *Drosophila* populations where dozens to hundreds of generations of selection have produced large differences in longevity and life history traits. Our study also represents a more nuanced investigation of the physiological mechanisms driving observed differences than previous studies which have been largely limited to assaying broad differences in total lipid and glycogen contents.

Metabolomic studies characterize differences in the prevalence of small-molecule metabolites in biological systems along some axis of interest. These collections of metabolites are viewed as one of the intermediary layers between the genome and expressed phenotypes, and thus represent a key component to understanding how phenotypes are determined. This general approach is already being used to explore the mechanisms driving physiological declines associated with advanced age ^30^, and *Drosophila-*based studies are already part of this effort ^31^. For instance, Hoffman et al. 2014 ^32^ analyzed changes in metabolomic profiles over time in different genetic backgrounds to identify aging-associated metabolic pathways. Taking a different approach, Laye et al. 2015^33^ examined how dietary restriction, a proven intervention strategy for extending lifespan in fruit flies, alters the metabolome to slow aging-related physiological declines. Here, we built on these efforts using an evolution-based approach where, like in the E&R studies described above, we used selection and replicate populations to amplify signals and identify relationships between metabolomic profiles and different aging patterns.

In this study, we investigated differences in metabolomic profiles among three groups of experimentally evolved *D. melanogaster* populations: populations where selection for rapid development has resulted in reduced longevity (A), and where selection for delayed reproduction (C) and starvation resistance (S) has led to increased longevity. In the past, the C populations have been used as controls for both the A and S populations. Studies comparing the C and A populations focused on the evolution and genetics of developmental rates, reproductive rates, and longevity ^9,29^, while work with the C and S populations focused more on understanding physiological changes associated with starvation resistance ^15^. However, with respect to longevity, we find similar mean lifespans in the C and S populations while the A populations tend to die at much younger ages (Figure 1A). Additionally, all populations are ultimately derived from a single ancestral population, differences between groups are known to be the result of selection on standing genetic variation ^15,26^, and populations cluster genetically based on treatment (Figure 1B). As such, comparisons across the three groups have the potential to isolate and identify mechanisms specifically associated with longevity (i.e. elements that overlap between the C and S populations but differ in A). Using metabolomic data from these populations we aimed to: 1) identify metabolites associated with longevity differences based on patterns of overlap and differentiation between groups, and 2) identify genetic mechanisms underlying based on statistical associations between patterns of genomic and metabolomic differentiation.

**Figure 1.**
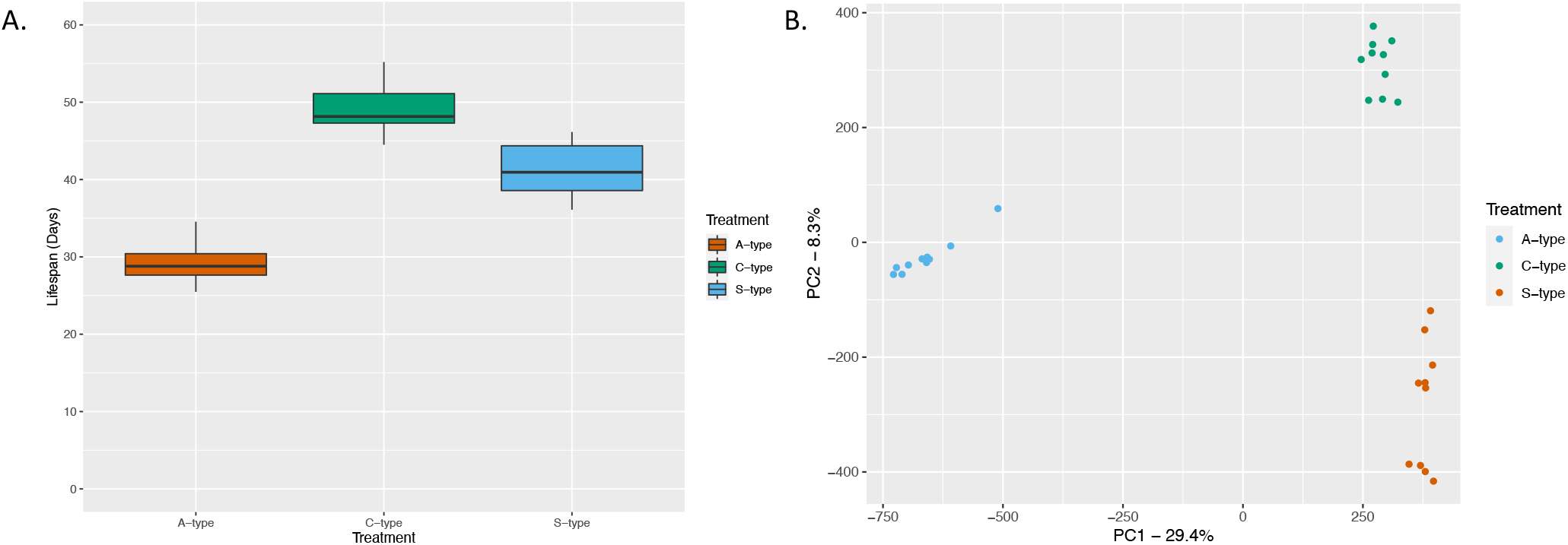
Lifespan comparison for the A, C, and S populations (A), and PCA plot showing how populations cluster based differences and similarities in SNP frequencies across the genome (B). Plots were generated using data from Burke et al. (2016), Graves et al. (2017), and Kezos et al. (2019).

## Materials and Methods

### Experimental populations

This study involved 30 large outbred, experimentally evolved *D. melanogaster* populations evenly split between three treatments (A, C, and S). All populations trace back to a single ancestral laboratory population created by Phillip Ives by sampling 200 females from a wild population in Amherst, Massachusetts ^34^. In 1980, samples were taken from this now laboratory adapted population by Michael Rose and Brian Charlesworth for use in selection experiments to study life history evolution ^7,35,36^. This has since expanded to an experimental radiation consisting of dozens of populations split across a wide array of selection treatments maintained in the Rose Lab at UC Irvine. The A and C populations are described in depth in Burke et al. ^9^ and Graves et al. ^26^. Briefly, selection in these treatments is defined by optimization around different reproductive windows. The A populations are maintained on a discrete 10-day generation cycle, while the C populations are maintained on a 28-day cycle. Over generations, this has resulted in accelerated development and reduced longevity in the A populations relative to the C populations (see Burke et al. 2016 for in-depth phenotypic characterizations^9^). Populations are also maintained at a minimum census size of ~2000 individuals to avoid inbreeding. Both the A and C groups consist of two sub-groups: ACO_1-5_/AO_1-5_ and CO_1-5_/nCO_1-5_, respectively. The primary difference between these groups is the number of generations under selection; AO and nCO were more recently derived than the ACO and CO populations. At the time of sampling for this project, the ACO and AO populations had ~1000 and ~490 generations under selection respectively, while the CO and nCO populations have ~415 and ~160 generations. The broad differences between the A and C groups are due to the fact generation length is shorter in the former than the latter (10 vs 28 days). Within a group, previous efforts previous efforts have found extreme phenotypic ^9^, genomic ^26^, and transcriptomic ^29^ convergence between subgroups within a given treatment (e.g. there are no statistically significant differences between the ACO and AO populations). As such, we now simply recognize them as the A and C populations.

Unlike the A and C populations where selection is defined by shifting reproductive windows, the S populations have been subjected to selection for increased starvation resistance. A full description of the S populations and how they were created can be found in Kezos et al. ^15^. Briefly, the 10 S populations (SCO_1-5a_ and SCO_1-5b_) were derived from the CO_1-5_ populations with two replicates derived from each CO population (e.g., SCO_1a_ and SCO_1b_ were derived from CO_1_). In terms starvation selection protocols, after 2 weeks of development from eggs to adults in vials, approximately ~3000 flies are collected from each population and split between Plexiglas cages. This two-cage approach is used to maintain high census sizes through selection. At this point, flies are fed an agar media prepared with cooked bananas, corn syrup, yeast, and barley malt (see Rose et al. 2004 for exact recipe)^10^. On day 17 from egg, Petri dishes with banana agar media are removed from cages and replaced with dishes containing media made with only agar and water (i.e. a nonnutritive media). This starvation condition is maintained until 75–80% of flies have died. At this point, the two cages set up for each replicate are condensed into one cage. Flies are then provided with banana agar media for days at which points eggs are collected to start the next generation. From generation 50 onwards, starvation conditions typically needed to be maintained for ~10 days to reach the 75–80% mortality threshold. At the time of this study the S populations had experienced ~160 generations of selection.

Lastly, with 10 replicate populations per treatment group and dozens to hundreds of generations under selection, we would expect this study to have a great deal of statistical power in a traditional E&R context ^37^. And while this type of theoretical power analysis has not been done for studies combining experimental evolution and metabolomics, it stands to reason that the statistical power stemming from population level replication and using evolution to create patterns of differentiation should be similarly useful when seeking to identify candidate metabolites.

### Collecting flies for metabolite extraction

To collect samples for metabolomic characterization, we used the same sampling procedures described in previous genomic studies ^15,26^. Briefly, cohorts were derived from each replicate population and reared on a 14-day culture cycle for two generation to reduce the impact of maternal effects. Cohorts descended from the second cycle were sampled for metabolite extraction. At creation, each cohort consisted of ~1500 flies in a Plexiglas cage and were fed the standard banana-based media with food being changed every two days. On day 21 from egg, 150 females were collected from each cohort, flash frozen in liquid nitrogen, and stored at −80°C until it was time for extraction.

The decision to collect samples on day 21 for metabolomic characterization was motivated by the findings of Barter et al. ^29^. Here the authors compared gene expression in the A and C populations at two time points, day 21 and day 14, and found that differences much greater differences between the groups at the later timepoint. This was ultimately attributed to the fact that in demographic terms, the populations are not yet aging at day 14 (i.e. there is no acceleration in mortality during this period). However, by day 21, the A populations are clearly aging by this definition while the C populations are not^9^. And this is ultimately being reflected in patterns of differential gene expression at different life stages with there being greater power to identify aging related genes when one group is in its aging phase and the other is not. Given this, it stands to reason that comparing metabolomic differences at day 21across between treatments should similarly allow for more powerful identification of metabolic pathways underlying different rates of aging between the groups.

### Metabolomic characterization

#### Gas chromatography-mass spectrometry (GC-MS)

For metabolite extraction, samples were extracted in −80°C 2:2:1 methanol/acetonitrile/water that contained a mixture of nine internal standards (d_4_-Citric Acid, ^13^C_5_-Glutamine, ^13^C_5_-Glutamic Acid, ^13^C_6_-Lysine, ^13^C_5_-Methionine, ^13^C_3_-Serine, d_4_-Succinic Acid, ^13^C_11_-Tryptophan, d_8_-Valine; Cambridge Isotope Laboratories) at a concentration of 1 μg/ml each. The ratio of extraction solvent to sample volume was 18:1. Fly tissue samples were lyophilized overnight prior to extraction. After the addition of extraction buffer, fly tissues were homogenized using a ceramic bead mill homogenizer. The samples were then incubated at −20°C for 1 hour followed by a 10-minute centrifugation at maximum speed. Supernatants were transferred to fresh tubes. Pooled quality control (QC) samples were prepared by adding an equal volume of each sample to a fresh 1.5-ml microcentrifuge tube. Processing blanks were utilized by adding extraction solvent to microcentrifuge tubes. Samples, pooled QCs, and processing blanks were evaporated using a speed-vac, and the resulting dried extracts were derivatized using methyoxyamine hydrochloride (MOX) and N,O-Bis(trimethylsilyl)trifluoroacetamide (TMS) [both purchased from Sigma]. Briefly, dried extracts were reconstituted in 30 μl of 11.4 mg/ml MOC in anhydrous pyridine (VWR), vortexed for 10 min, and heated for 1 hour at 60°C. Next, 20 μl TMS was added to each sample, and samples were vortexed for 1 minute before heating for 30 minutes at 60°C. The derivatized samples, blanks, and pooled QCs were then immediately analyzed using GC-MS.

GC chromatographic separation was conducted on a Thermo Trace 1300 GC with a TraceGold TG-5SilMS column (0.25 um film thickness; 0.25mm ID; 30 m in length). The injection volume of 1 μl was used for all samples, blanks, and QCs. The GC was operated in split mode with the following settings: 20:1 split ratio, split flow: 24 μl/min, purge flow: 5 mL/min, Carrier mode: Constant Flow, Carrier flow rate: 1.2 mL/min. The GC inlet temperature was 250°C. The GC oven temperature gradient was as follows: 80°C for 3 minutes, ramped at 20°C/minute to a maximum temperature of 280°C, which was held for 8 minutes. The injection syringe was washed 3 times with pyridine between each sample. Metabolites were detected using a Thermo ISQ single quadrupole mass spectrometer, and data were acquired from 3.90 to 21.00 minutes in EI mode (70eV) by single-ion monitoring (SIM). Metabolite profiling data were analyzed using TraceFinder 4.1 utilizing standard verified peaks and retention times.

We used TraceFinder 4.1 to identify metabolites in extracted samples, blanks, and QCs by comparing sample metabolite peaks against an in-house library of standards prepared by processing and analyzing authentic standards via the method described above. We created a database of retention times and three fragment ions for each metabolite standard: a target peak/ion and two confirming peaks/ions. When running biological samples, we identified metabolites that not only matched with the known retention times of the authentic standard but also with its target and confirming peaks. TraceFinder was also used for GC-MS peak integration to obtain peak areas for each metabolite. After TraceFinder analysis, we corrected for instrument drift over time using local regression analysis as described by Li et al. ^38^ We use the pooled QC samples, which were run in duplicate at the beginning and end of the GC-MS run for this purpose. The data are then normalized to an internal standard to control for extraction, derivatization, and/or loading effects.

#### Liquid chromatography-mass spectrometry (LC-MS)

Fly samples were dried to make extracts. Dried extracts were reconstituted in 40 μL acetonitrile/water (1:1 v/v), vortexed well, and transferred to LC-MS autosampler vials for analysis. LC-MS data were acquired on a Thermo Q Exactive hybrid quadrupole Orbitrap mass spectrometer with a Vanquish Flex UHPLC system or Vanquish Horizon UHPLC system. Notably, the LC column used was a Millipore SeQuant ZIC-pHILIC (2.1 × 150 mm, 5-μm particle size) with a ZIC-pHILIC guard column (20 × 2.1 mm). The injection volume was 2 μL. The mobile phase was composed of solvent A (20 mM ammonium carbonate [(NH4)2CO3] and 0.1% Ammonium Hydroxide [NH4OH]) and solvent B (Acetonitrile). The mobile phase gradient started at 80% solvent B, decreased to 20% solvent B over 20 minutes, returned to 80% solvent B in 0.5 minutes, and was held at 80% for 7 minutes. (PMID: 28388410). The method was run at a flow rate of 0.150 mL/min. Subsequently, the mass spectrometer was operated in full-scan, polarity-switching mode from 1 to 20 minutes, with the spray voltage set to 3.0 kV, the heated capillary held at 275 °C, and the HESI probe held at 350 °C. The sheath gas flow was set to 40 units, the auxiliary gas flow was set to 15 units, and the sweep gas flow was set to 1 unit. MS data acquisition was performed in a range of m/z 70–1,000, with the resolution set at 70,000, the AGC target at 1 × 106, and the maximum injection time at 200 ms ^38^

For data analysis, acquired LC-MS data were processed by Thermo Scientific TraceFinder 4.1 software, and metabolites were identified based on the University of Iowa Metabolomics Core facility standard-confirmed, in house library. NOREVA was used for signal drift correction ^26,38^. Data were normalized to total ion signals, and MetaboAnalyst 4.0 was used for further statistical processing and visualization^39,40^.

#### Analyzing Metabolomic Data

We conducted analyses using the MetaboAnalyst 4.0 webservice (http://www.metaboanalyst.ca). Metaboanalyst is a module that uses both statistical and machine learning methods to provide visualization for classifying our data into groups. This program was utilized for heat mapping, enrichment pathways, pathway analysis, and statistical analysis. Unless otherwise noted, all data are reported as mean ± SD. One-way ANOVA followed by Tukey multiple comparison test was utilized. A probability value of P ≤ 0.05 was considered significantly different. Statistical calculations were performed using the GraphPad Prism software (La Jolla, CA, USA).

### Linking genome to metabolome

Here the goal was to identify patterns of SNP differentiation between the A, C, and S populations that best predict key patterns of metabolomic differentiation. For this analysis, efforts were focused on significantly differentiated metabolites associated with the top ten enriched pathways from our GC-MS and LC-MS metabolomic profiling (see Supplementary Table 1 for list of metabolites, and Figures 2C and 3C for enriched pathways). SNP data came from previously published pool-seq DNA data from the (see Graves et al. 2017^26^ for DNA extraction details for the A and C populations, and Kezos et al. 2019^15^ for the S populations). Genomic data was reprocessed to since the original studies used different version of the *D. melanogaster* reference genome. However, processing steps were otherwise the same. Briefly, fastq files corresponding to each population were mapped to the *D. melanogaster* reference genome (version 6.14) with BWA^41^ using bwa mem with default setting, and SAMtools ^41^ was used to convert the resulting SAM files to BAM files, remove potential PCR duplicates, and merge all BAM files into a single mpileup. PoPoolation2 ^37^ was used to convert this mpileup file to a simplified file format that contains counts for all bases in the reference genome and for all populations being analyzed. RepeatMasker (http://www.repeatmasker.org) and PoPoolation2 were then used to identify and remove highly repetitive genomic regions where proper read mapping is difficult. Lastly, SNPs were called based on the following criteria: minimum coverage of 20X and maximum of 200X in each population, and a combined minor allele frequency of 2% across all populations. This resulted in a SNP table with ~781K sites.

**Figure 2.**
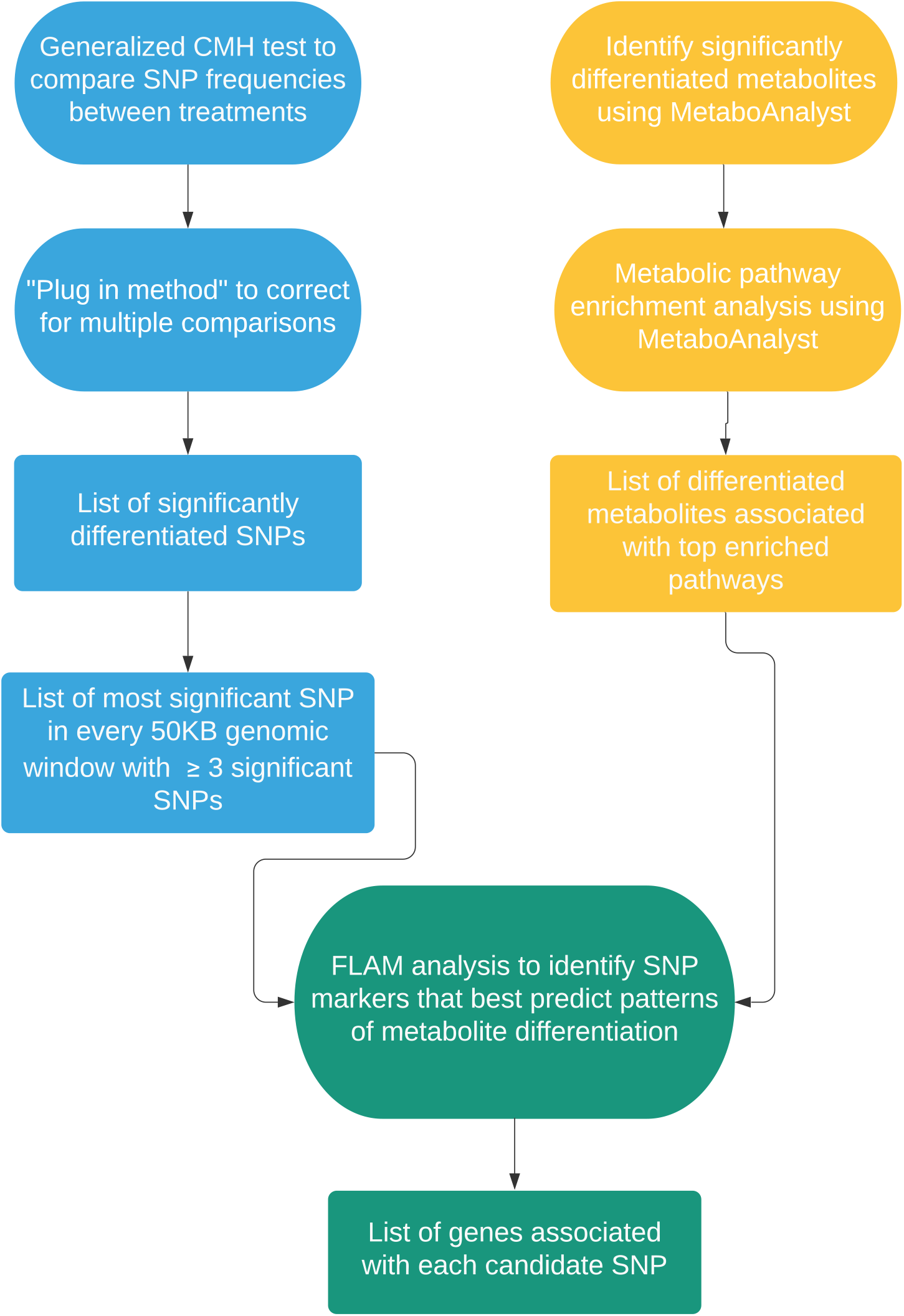
A flowchart outlining methods to identify candidate genes associated with differences in candidate metabolites.

**Figure 3.**
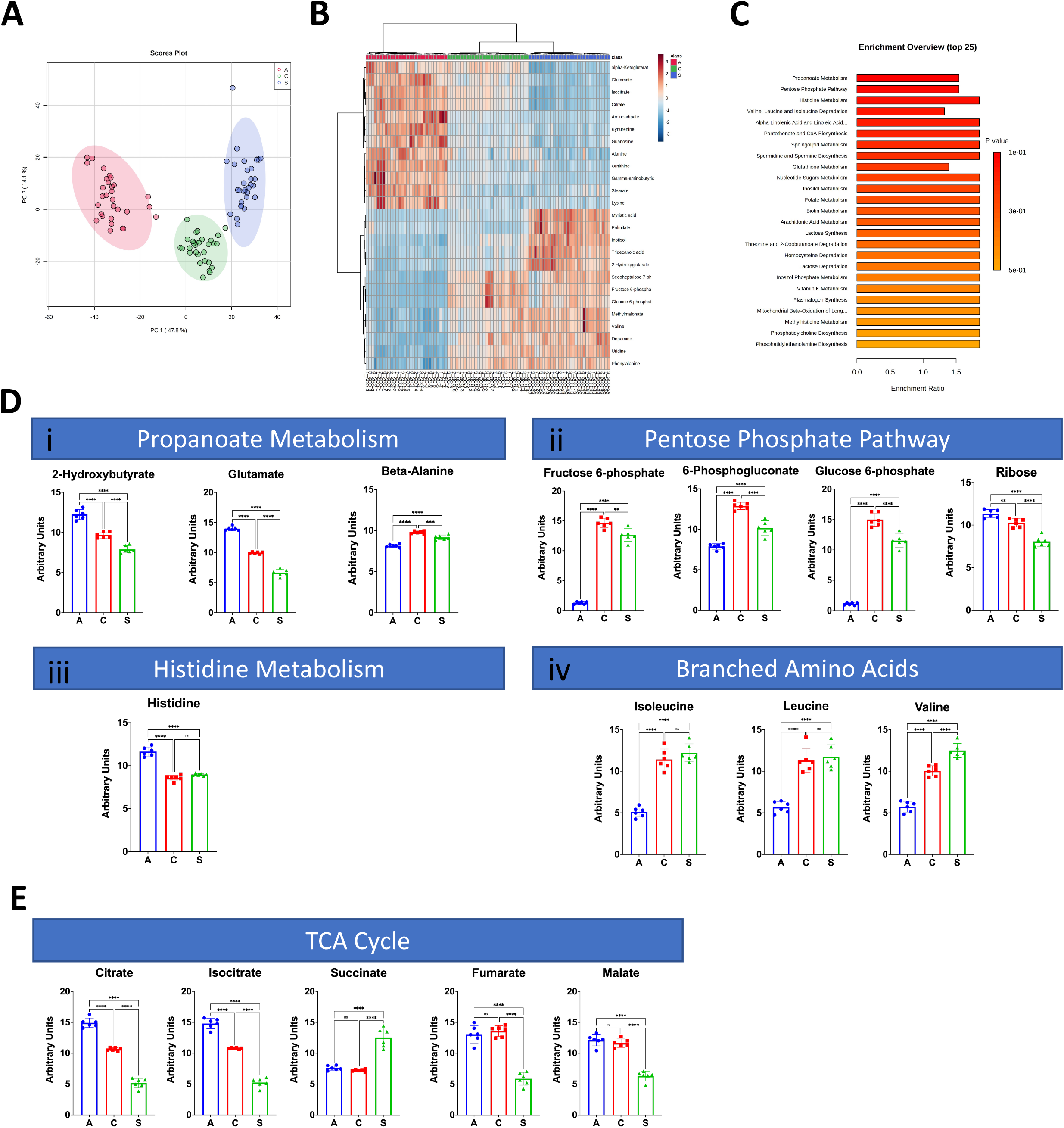
Figure 3: GC-MS Metabolomic Profiling of A, C, and S populations. PCA plot of A, C, and S metabolite profiles (A). Heatmap comparison of top 25 changed low energy state metabolites among A, C, and S populations (B). Top 25 enriched metabolic pathways (C). Quantification of metabolites from the top 4 enriched metabolic pathways among A-, C-, and S populations (D). Key components of the TCA cycle (E) were among the top 25 changed metabolites. Mean ± SD; n = 6; *p-value ≤ 0.05.

Figure 2. outlines the analytic approach used to link patterns of metabolomic and genomic differentiation. First, the Generalized Cochran-Mantel-Haenszel (CMH) test ^42^ was used to identify SNPs that were significantly differentiated between the A-, C-, and S populations. This was done in R using the mantelhaentest function ^43^ tests were performed for each SNP in the data set, correcting for multiple comparisons was essential. To do this, the “plug in method” (Hastie et al. 2009) was used. Briefly, suppose we have *M* total hypothesis tests, and we let *V* be the number of false positives and S be the number of true positives, then the false discovery rate (FDR) is defined as *V/(V+S*). A critical test statistic, C, can be chosen and the plug-in method computes the FDR for that critical point*. V+S* is estimated to simply be the total number of test statistics from the M hypothesis tests that exceed C. V can in turn be estimated through permutation. Here that was done by essentially shuffling population labels then performing the CMH test at each polymorphic site in the permuted data set. The number of significant test statistics greater than all values of *C* was then recorded. This was repeated 100 times to ensure accurate estimates of *V* for values of *C*. Once *V* and *V+S* were estimated, the FDR was calculated for each value of *C*. Using these results, we calibrated to a conservative FDR of 0.005 (i.e. significance threshold is the value of *C* that gives an FDR of 0.005).

After identifying SNPs that were differentiated across our treatments, a statistical learning approach called the “fused lasso additive model” or “FLAM” ^44^ was used to determine which of the genomic regions these SNPs represent best predict patterns of differentiation for our top candidate metabolites (Note: this was done on a per metabolite basis). Here the assumption is that these genomic regions are the ones most to be casually linked to relevant patterns of metabolite differentiation. Due to genetic linkage, it is not necessary to consider individual significant SNP. Instead, a list of the most significant SNP for every 50KB genomic window and these markers were used in the FLAM analysis. This mimics the implementation of FLAM as described in Muller et al^23^ where the approach was validated using simulated and real data sets. It is also worth noting that this implementation strategy differs from the original Petersen et al^44^ implementation in one major way. As described in Mueller et al.^23^, a single run of FLAM is limited to finding *N* casual SNPs, where *N* is the number of populations in the study. As a result, the order of potential predictor variables can impact results. Here and in Muller et al. ^23^ this is accounted for using a permutation procedure where each FLAM analysis is run multiple times and the order of potential predictor variables is randomly shuffled. The final list of “best predictors” consists of genetic loci that occur at commonly identified across permutations. In this study, a total of 100 permutations were run for each metabolite, and the final list of best predictors consisted of loci that showed up in at least 50% of permutations of a given metabolite. Lastly, based on past validation, FLAM itself is particularly well suited for the task of linking genomic and metabolomic differentiation in an E&R context. A major consideration in these studies is the need to distinguish between parallel differences with some underlying relationship with the phenotype of interest, and differences simply due to chance. Muller et al. ^23^ suggests that FLAM has the power to distinguish between these two groups based on subtle differences among replicate populations within a treatment, which is not always possible with standard linear model approaches.

After determining which genomic regions best predict observed differences in candidate metabolites, a list of genes associated with each region was generated. Given that predictors are markers representing genomic regions, it cannot simply be assumed that the SNP markers are themselves casual. However, given that the SNPs used in the FLAM analyses are the most significant SNPs in their respective genomic region and the and the level of replication featured in this study, we would expect them to be relatively close to the true causative sites given theoretical work on the power of E&R studies to localize candidate genes ^45^. Candidate genes were ultimate defined as those in 5KB windows around each SNP marker with this in mind. After generating a list of candidate genes associated with each candidate metabolite, Metascape ^46^ was used to perform gene ontology (GO) term enrichment analysis, protein network analysis, and Molecular Complex Detection (MCODE) Component analysis. All analyses were run using default settings. Cytoscape ^47^ was used to visualize results.

## Results

### Metabolomic results

To identify metabolic changes associated with life history, a detailed characterization of whole-body metabolism was undertaken in A, C, and S populations using both GC-MS and LC-MS.

#### Characterization of GC-MS Metabolomic Profiling of A, C, and S populations

GC-MS analysis returned 104 metabolites, 48 of which were significantly changed between populations. Principal component analysis (PCA) of metabolites revealed three distinct metabolic populations, with PC1 and PC2 explaining 47.8% and 14.1% of the variability, respectively (Figure 3A). The S and C populations were similar to one another and distant from A along PC1. Visualization of the top 25 most significant metabolites revealed a distinct inverse relationship between A and S metabolomic profiles, with the C profile appearing intermediate (Figure 3B). Enrichment analysis revealed propanoate metabolism, the pentose phosphate pathway, histidine metabolism, and valine, leucine, and isoleucine degradation as the top metabolic hits associated with life history (Figure 3C). Closer inspection of propanoate metabolism revealed that C and S populations had decreased 2-hydroxybuterate and glutamate, and increased beta-alanine compared to A (Figure 3Di). In the pentose phosphate pathway, fructose 6-phosphate, 6-phosphogluconate, and glucose 6-phosphate were elevated in the C and S populations compared to A (Figure 3Dii). Histidine was decreased in C and S populations compared to A (Figure Diii). Isoleucine, leucine, and valine were elevated in C and S compared to A (Figure Div). Components of the tricarboxylic acid (TCA) cycle, a critical pathway for aerobic respiration, were also significantly associated with life history (Figure 3E). Citrate and isocitrate levels decreased in a stair-step manner from A to S. Interestingly, succinate, fumarate, and malate levels were not significantly different between A and C populations. In the S population, succinate levels were higher while fumarate and malate levels were lower than A and C populations.

#### Characterization of LC-MS Metabolomic Profiling of A, C, and S populations

To broaden our metabolic characterization of A, C, and S populations, we also used liquid chromatography mass spectrometry (LC-MS) for better characterization of non-volatile, thermally unstable, and higher molecular weight species. LC-MS analysis returned 38 metabolites, of which were significantly changed between populations. PCA revealed a much greater degree of separation among these metabolites than that uncovered by GC-MS. PC1 and PC2 explained 51.2% and 16.6% of variability, respectively (Figure 4A). There was less distinction between C and S populations along PC1, with most variability between the two explained by PC2. The A type population remained separate and distinct (Figure 4A). Accordingly, visualization of the top 25 significant metabolites revealed similar profiles among C and S populations and both appeared to have an inverse relationship to that of the A (Figure 4B). Fatty acid metabolism, beta-oxidation of long and short chain fatty acids, and methionine metabolism were the top enriched metabolic processes identified (Figure 4C). Short and long-chain species of carnitine were significantly lower in the A population compared to the C and S populations (Figure 4D). Similarly, methionine levels were significantly lower in the A population compared to C and S populations (Figure 4D). Key metabolic co-factors, FAD, NAD+, and NADP+, were also identified among the top 25 significant metabolites (Figure 4E). FAD and NAD+ levels among the C and S populations were elevated compared to A, while NADP+ levels were decreased.

**Figure 4:**
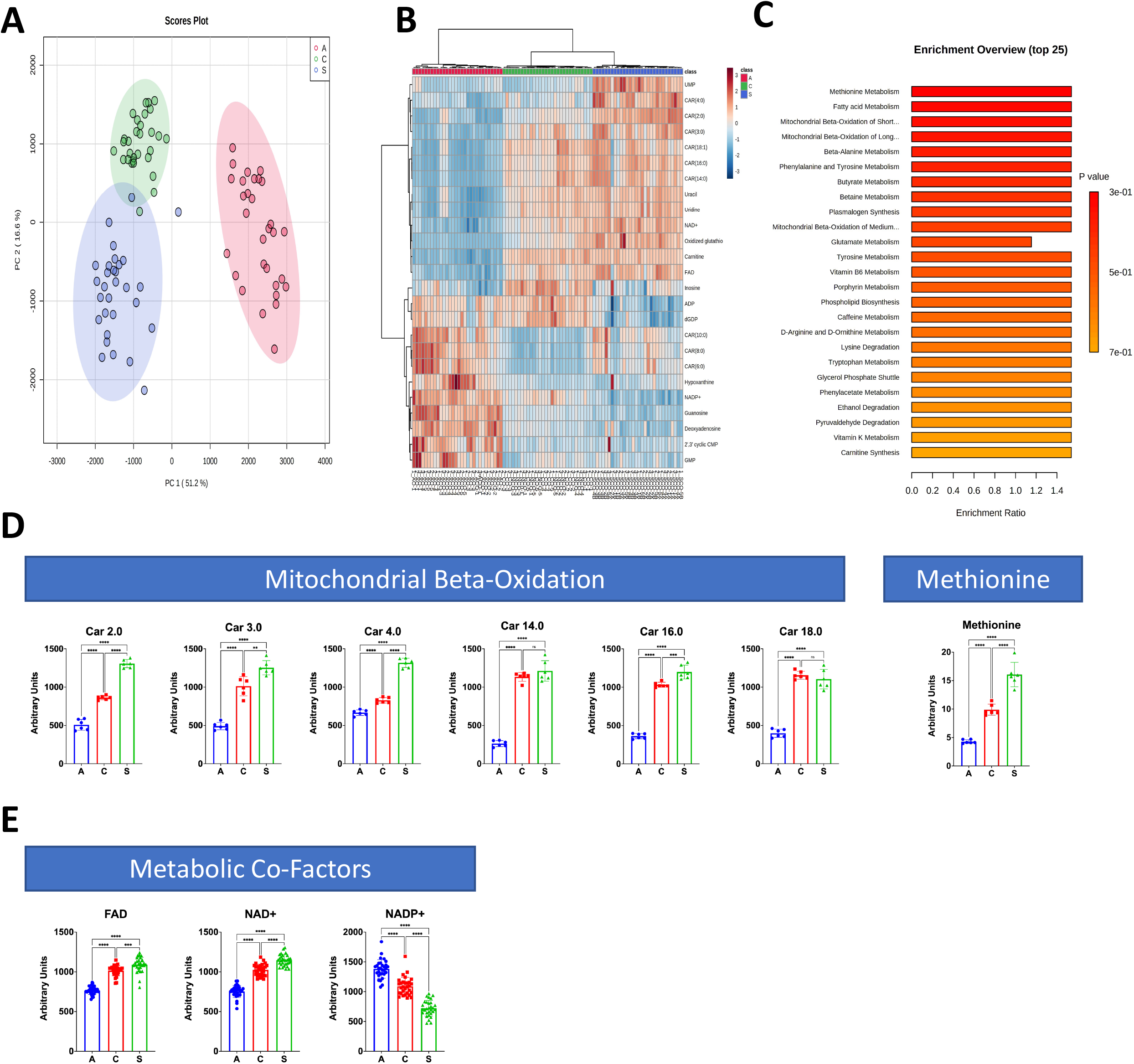
LC-MS Metabolomic Profiling of A, C, and S populations. PCA plot of A, C, and S populations high energy state metabolite profiles (A). Heatmap comparison of top 25 changed high energy state metabolites among A, C, and S populations (B). Top 25 enriched metabolic pathways (C). Quantification of metabolites in select enriched pathways from A, C, and S populations (D). Select important metabolic co-factors (E). Mean ± SD; n = 6 runs, 5 flies/run/group; *p-value ≤ 0.05.

### Genome to Metabolome

Linking patterns of SNP variation between the A, C, and S populations to top candidate aging metabolites was a multistep process (see Figure 2 for an overview). Conventional statistical methods were first used to identify significantly differentiated SNPs between the three groups. This resulted in a list of ~76K SNPs. However, as many of SNPs are likely neutral variants linked to causative sites, this list was reduced to the most significant SNP in every 50KB genomic window with at least 3 significant SNPs which yielded 1827 SNP markers. FLAM, a statistical learning approach, was then used to determine which markers best predicted patterns of differentiation across the A, C, and S populations for each of candidate metabolite. This resulted in a total of 221unique SNP markers with values ranging for 3 to 36 top predictors per metabolite (Supplementary Table 1). These top candidates were found across all major chromosome arms, and there was no clear relationship with chromosome length and number of candidates (Supplementary Figure 1 and Supplementary Table 2). Looking at the mean SNP frequencies for each marker in each group of populations, we typically find the expected pattern where C and S are similar in value and different from A (Supplementary Figure 1). Next, unique markers identified by FLAM were converted to a gene list based on genes present in 5-kb windows around each marker. This resulted in a list of 494 candidate genes (Supplementary Table 3 for a list of all genes and associated GO terms).

Metascape, which clusters terms into groups based on similarity, was used to identify enriched GO terms and pathways given our list of candidate genes. For enriched GO term clusters, there was a common trend of terms associated with development and growth, especially with regards to the nervous system (Figure 5A and Supplementary Table 4 for more details). We also identified several GO terms associated with metabolism (e.g. regulation of RNA metabolic processes and glycolytic processes). Examining the relationships between these clusters, we found a pattern where nodes representing development, growth, and nervous system-related clusters largely grouped together while terms related to metabolic processes were more distal or even unconnected like in the case of the glycolytic processes cluster (Figure 5B). In addition to GO term enrichment, we also used Metascape to perform Protein Network and MCODE Component analyses. Here we again found similar trends with enriched terms from the Protein Network analysis being associated with neurological development, and terms from the MCODE component analysis being associated with metabolic processes (Table 1).

**Figure 5.**
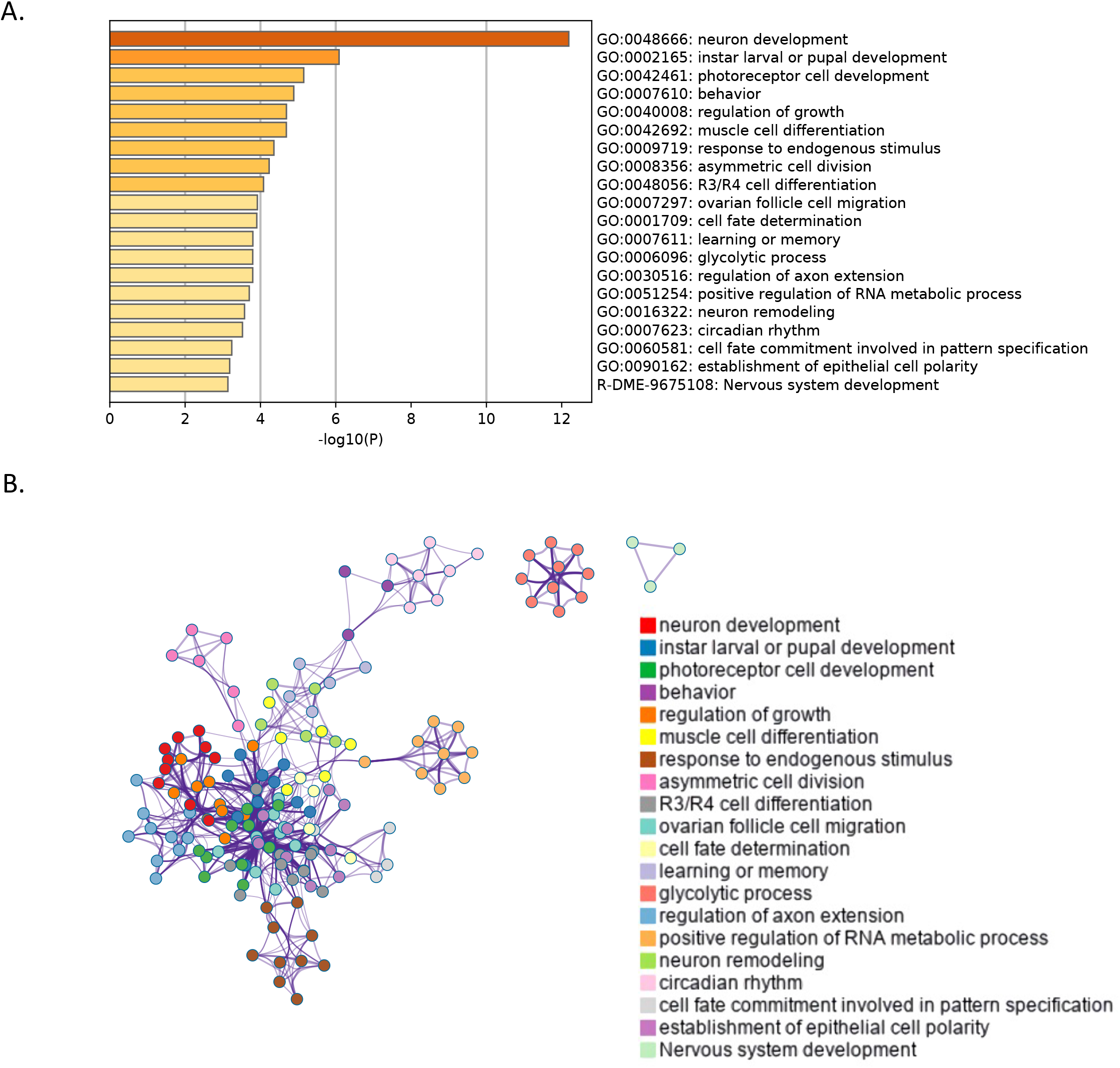
Heatmap of top 20 enriched GO clusters (A), and a network showing relationships between clusters (B).

**Table 1.**
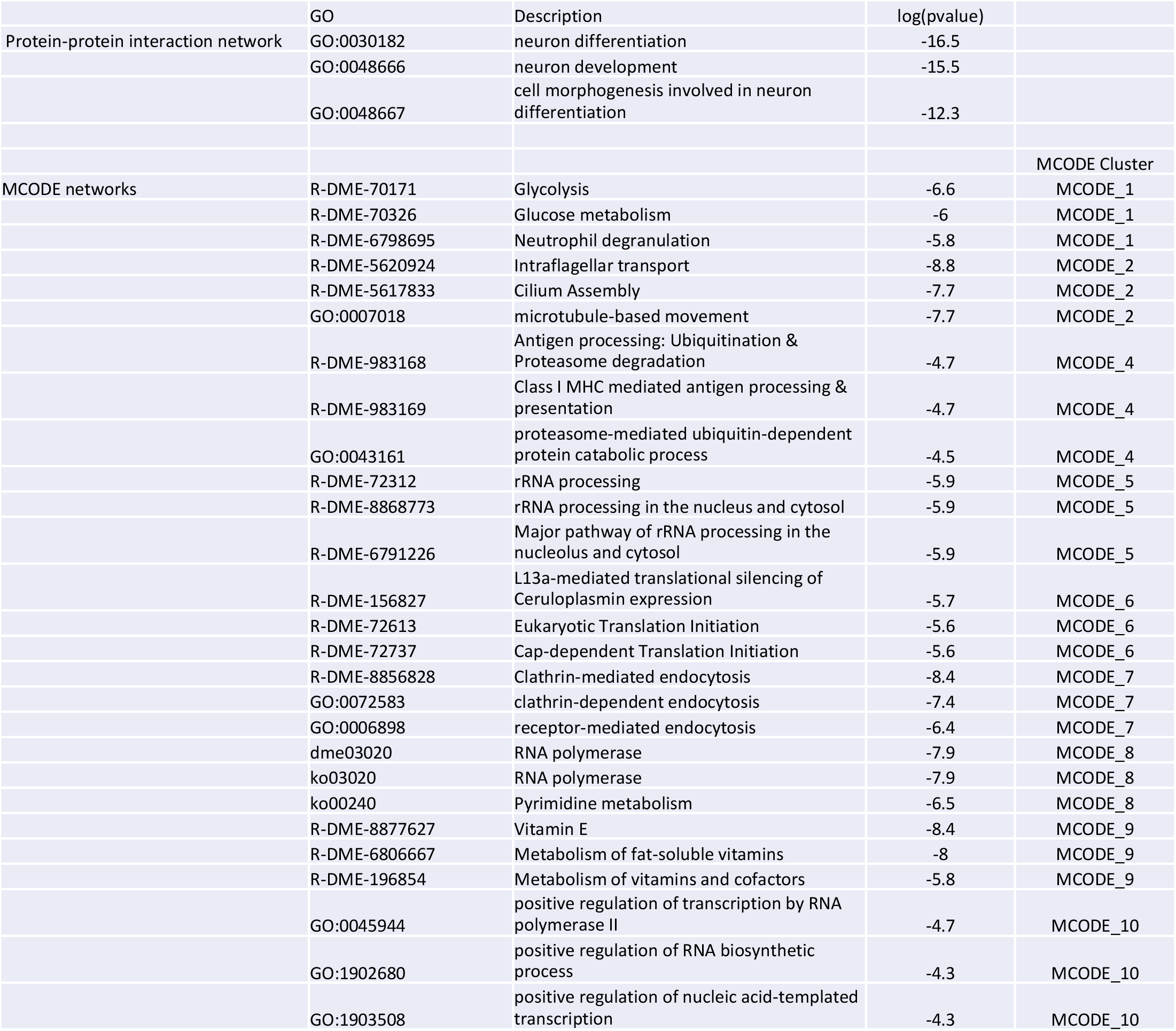
Results of protein-protein enrichment analysis and MCODE component analysis.

## Discussion

### Metabolomic characterization

One of the primary objectives of this study was to identify metabolomic profiles associated with longevity differences by incorporating metabolomic characterization into the E&R framework (GC-MS: Figure 3 and LC-MS: Figure 4). GC-MS is the gold standard for identifying compounds that are volatile and thermostable. At the same time, LC-MS is more adept at identifying non-volatile, thermally unstable, and more significant compounds ^48^. Metabolites returned from GC-MS analysis were, to a degree, able to differentiate between A and S populations. For most metabolites returned, these two groups were inversely related, suggesting these metabolites are essential for understanding the driving mechanisms for starvation-resistant longevity (Figure 3B). However, care should be taken in the selectivity of these metabolites in association with longevity as the metabolic profile of the long-lived C population oscillated between that of the A and S populations. For example, 60% of TCA metabolites in the C population correlated with that of the short-lived A population (Figure 3E), whereas 67% of branched amino acids correlated with that of the long-lived S population (Figure 3Di-Figure 3Dv).

Deepening the metabolic characterization through LC-MS, we identified profiles that better-reflected lifespan (Figure 4). Metabolites returned from this analysis showed clustering of both long-lived populations (C and S) and greater separation from the short-lived population (A) (Figure 4A). Of the top 25 significant metabolites identified, ~70% correlated between C and S populations, suggesting that further analysis and exploration of these hits could yield greater insights into mechanisms underlying longevity differences in this system. For example, NAD^+^ is an abundant coenzyme that acts as the oxidizing-reducing agent inside the cell ^30,49,50^. NAD^+^ plays a necessary role for hydrogen transfer in redox reactions through accepting hydride from metabolic processes, including glycolysis, the TCA cycle, and fatty acid oxidation (FAO), to form NADH. NADH serves as a central hydride donor to ATP synthesis through mitochondrial oxidative phosphorylation, leading to the generation of reactive oxygen species. Studies have shown that NAD^+^ concentrations decline with age in worms, flies, mice, and humans ^51–58^. For instance, decreasing NAD^+^ levels in *C. elegans* results in a reduction in lifespan ^59–61^. The depletion of NAD^+^ is linked to mitochondrial dysfunction, mtDNA genomic instability, and deregulated nutrient sensing ^55,62,63^. This suggests that the maintenance of higher NAD^+^ levels is beneficial, potentially through maintenance of mitochondrial homeostasis.

NAD^+^ is also a co-substrate for numerous enzymes including sirtuins^64,65^, PARPs, CD38, CD157, CD73, and SARM1^66,67^. NAD^+^ serves as a required substrate for the deacetylase activity of the sirtuin family of proteins (SIRTs). Mitochondrial SIRTs, SIRT3, −4, and −5, are connected in metabolism, mitochondrial fidelity, and cell stress. SIRT3, SIRT4, and SIRT5 are found primarily located in the mitochondria, and are implicated in several of the principal processes of this organelle. SIRT3 has been the subject of serious investigation and is fundamentally a deacetylase estimated to function as a mitochondrial fidelity protein, with roles in mitochondrial substrate metabolism, protection against oxidative stress, and cell survival pathways ^68–70^. Less is known about the useful targets of SIRT4, which has deacetylase, ADP-ribosylase, and a newly defined lipoamidase function ^71–73^. SIRT5 modulates acyl modifications including succinylation, malonylation, and glutarylation in both mitochondrial and extra-mitochondrial compartments. However, the functional significance of SIRT5 in the regulation of many of its proposed target proteins remains to be discovered^74–76^. Sustained mitochondrial stress leads to mitochondrial unfolded protein stress response stress and ER stress^77–81^. Data suggest that the short-lived flies would have accelerated mtUPR and sustained levels of ER stress, whereas the C and S, longer lived groups would have a blunted effect. Our findings here suggest a potential elevation in NAD^+^ consumption or decrease in synthesis in the A population compared to C and/or S (Figure 4E). Paired with a decreased presence of beta oxidation products and elevated TCA cycle substrates, our findings may suggest decreased mitochondrial function in the A population (Figure 3E and Figure 4). Thus, decreased NAD^+^ availability may be a contributing and modifiable factor in age-related diseases. The mechanisms controlling its levels in aging, however, remain incompletely understood.

NAD^+^, through SIRT1, can regulate the mTOR pathway ^76^. Mechanistic target of rapamycin (mTOR), or TOR in *Drosophila*, is a key regulator of metabolism. mTOR is composed of two distinct kinase complexes, mTOR complex 1 (mTORC1) and 2 (mTORC2), which are characterized by the signature components Raptor and Rictor, respectively^76,82–84^. Branched chain amino acids (BCAA), including leucine, isoleucine, and valine, play a crucial role in the activation of the TOR pathway^85^. Mitochondrial stress can be regulated by Tor through activation of ATF4 to induce the integrated stress response and activation of GCN2 to increase amino acid availability ^86–94^, which we believe to be a protective response. We hypothesize that the lower BCAA levels in the A population (Figure 3Di) indicate that they are not able to effectively activate this pathway. We believe the elevated BCAAs in the long-lived populations (Figure 3iv) suggest greater TOR activation may be associated with longevity ^95^. suggests that the A populations may have reduced levels of ATF4 activation which inhibits the cells’ ability to regulate stress responses and cellular homeostatic process (Figure 3iv). Equally, under physiological glucose concentrations, mTORC1 is stimulated and leads to a number of proteins and enzymes being altered. These altered proteins are involved in anabolic processes, while restricting the autophagic process. Conversely, when glucose levels are low, mTORC1 is inhibited, in turn leading to the repression of numerous anabolic processes, sparing ATP and antioxidants. However, in the S populations, there are high levels of glucose and high levels of amino acids (Figure 3 and Figure 4). Therefore, suggesting that prolonged activation of mTOR leads to beneficial autophagy.

Our conclusion that longevity differences in this system are tied to mitochondrial function and increased TCA cycle activity are supported by a number of metabolomic studies across different systems and species. For instance, results from work comparing metabolomic variation across eleven *Drosophila* species reports evidence that age and lifespan are associated with TCA cycle activity ^96^. Another phylogenetic based study comparing metabolomic and transcriptomic profiles across seventy-six wild yeast isolates also find associations between replicative lifespan and TCA cycle activity and mitochondrial function ^97^. Lastly, in a mouse model, metabolomic profiling of young and old individuals also points towards differences in mitochondrial function as a hallmark of aging ^98^. We believe this agreement across studies is such disparate systems greatly substantiates our interpretation of the mechanisms driving longevity differences between the A, C, and S populations.

### Genomics to metabolomics

In additional to characterizing metabolomic profiles, another major goal of this study was to explore the relationship between patterns of genetic differentiation and differences in candidate aging metabolites. In our two-step analysis, we sought to filter out uninformative sites and identify regions of the genome that best predict focal patterns of metabolite differentiation (Figure 2). This ultimately yielded several hundred candidate genes, which suggests the underlying genetic architecture for observed metabolomic differences is highly polygenic (Supplementary Table 3). This is in keeping with findings from experimental evolution studies in sexually reproducing eukaryotes and studies on the genetics of complex traits as a whole ^21,99,100^. However, it is worth noting that focusing on genomic regions that best predict candidate metabolite differentiation narrowed candidates to hundreds of genes from thousands if we had relied solely on SNP differentiation between groups. As such, there appears to be some real value in combining different types of omic data when attempting to parse the genetic architecture of complex phenotypes like longevity.

Although longevity is a major axis of differentiation between the A, C, and S populations and our metabolomic analysis was structured to identify longevity associated metabolites, we did not find significant enrichment for genes directedly related to aging looking across our candidate genes (Figure 5A and Table 1). The failure to find an overrepresentation of canonical aging genes is not uncommon in E&R studies focused on longevity ^28,29,101^. This is perhaps due to a lack of segregating genetic variation at related loci as Fabian et al.^101^ suggests, and relates to the context dependent and complex nature of genotype to phenotype map. That aside, looking across our list of candidates, we do indefinity three major canonical aging genes: *Tor*, *Atg7,* and *Kermit*. *Tor* and *Atg7* in particular have clear links to our metabolomic results; *Tor* through its many well defined aging related function ^102–104^ and *Atg7* though its effects on autophagic activity and relationship to mitochondrial function and turnover ^105–107^. *kermit*, while less obviously connected to our metabolomic results, is a gene required for proper locomotive activity that when inhibited or overexpressed has strong negative impacts on longevity and survival ^108,109^. So, while we do not observe an overrepresentation of canonical aging, the ones present seem likely to be biologically meaningful.

Given that our metabolomic results point to towards differences in mitochondrial function as a major force driving longevity differences between the A, C, and S populations, we queried our list of candidate genes for related term (see Supplementary Table 3 for genes and associated terms). As with aging, we do not find enrichment for genes associated with mitochondrial function. However, in addition to the previously mentioned *Atg7* and *Tor*, we find a number of genes directly or predicted to be associated with mitochondrial function (e.g. *ND-B14, COX6AL,* and *CG15386*) and maintenance (*mtDNA-helicase*, *mre11, CG11975,* and *larp).* We also find a number of genes directly or predicted to be associated with endoplasmic reticulum stress (CG9934, CG8405, S2P), which as explained above can also impact mitochondrial function. So, while we do not find clear overrepresentation, we nevertheless again find some concordance between specific candidate genes and our metabolomic characterization. And the lack of enrichment could once again be due to a general lack of relevant segregating genetic variation at loci related to mitochondrial function. Strong purifying selection at such loci should perhaps be expected given the importance of maintaining mitochondrial function.

Focusing on our enrichment results (Figure 5 and Table 1) and total list of candidate genes (Supplementary Table 3), we find clear overrepresentation of candidates associated with nervous system development and maintenance. With the exception of *Atg7* and *kermit*, the genes in these clusters are not explicitly defined as aging genes based on past studies. However, decades of research has established a clear link between longevity and nervous system development, regulation and function ^110–115^. As such, we can reasonably interpret our findings as evidence that longevity differences between the A, C, and S populations are at least in part due to genes impacting the nervous system and these effects are reflected in the populations’ metabolomic profiles. Also, while less pronounced, we find evidence of many candidate genes (Supplementary Table 3) and enriched clusters (Figure 5 and Table 1) associated with various forms of carbohydrate metabolism (e.g. glucose metabolism, glycolytic processes, amino sugar metabolic processes, and glutathione metabolic processes). This ties directly into the metabolomic results described above, and such process again have clear links to aging and longevity ^116,117^.

`Comparing our results to work done by` Barter et al. ^29^ which used the A and C populations to study the genetic basis of longevity and development yields mixed results. For instance, while there is still a major theme of development, it comes in primarily the form of somatic muscle development and ecdysteroid metabolic processes instead. Enrichment for genes associated ecdysteroid metabolic processes are particularly suggestive given that ecdysteroids play an important role in guiding developmental transitions in insects^118^. Presumably the lack of enrichment for such genes in this present study is due to our inclusion of the S populations to hone in specifically on longevity associated genes. To that point, looking at the results Kezos et al. ^15^ which used the C and S populations to study the genetic basis of starvation resistance, we primarily find enrichment for genes associated with metabolic processes and not development terms. `Here we speculate that these contrasts suggest that our three treatment approach and the incorporation of metabolomic data has indeed allowed us to better focus on longevity candidates`.

While there presently no other Drosophila E&R studies on aging incorporating metabolomics we can directly compare our results to, there are a number of “traditional” E&R we can use for points of comparison: Fabian et al.^101^ Carnes et al. ^25^, and Remolina et al. ^28^. These three studies all selected for longevity differences by manipulating reproductive timing in a similar fashion to what was done in the A and C populations. With regards to patterns of enrichment between studies, we do fine a great deal of concordance. For instance, while there are examples of individual canonical aging genes being identified in all of these studies, like us they do not find overrepresentation. All of these studies also report evidence for candidates associated with metabolism. Running their results through Metascape for a more direct comparison to our results we also see that Carnes et al.^25^ and Fabian et al.^101^ identify candidates enriched for genes associated with nervous system development and function (Supplementary Figure 2). Here it should be noted that the population studied by Carnes et al.^25^, while different than our own in terms of exact selective pressures and housed outside of the Rose Lab at UC Irvine for many generations, are ultimately derived from same ancestral population as the A, C, and S populations. However, this is not the case for the populations studied by Fabian et al.^101^. As such, this pattern cannot simply be dismissed as idiosyncratic to systems derived from the Ives’s domesticated laboratory population. In total, we would argue that these similarities between our studies further support the idea we are capturing true signals.

That being said, the overlap between the results of Carnes et al.^25^, Remolina et al. ^28^ and Fabian et al.^101^ and our study is not perfect. For instance, these studies all report that genes associated with immune function appear to be major drivers of differentiation between their populations. We do not find clear evidence of this in our study. Patterns of overlap between the lists of candidate genes in particular also complicate interpretation. Despite the overlap in enrichment for functional categories we do observer, actual gene overlap is modest (Figure 6, note this figure was made using the UpSetR^119^ package in R). Pairwise comparisons typically yield a few dozen shared genes and these numbers drop further when we compare across more than two studies. In fact, only 2 genes, *cac* and *shakB*, are shared across all three studies and our own. We believe this discordance between studies speaks to the complex and context dependent genetics of traits like longevity. It also supports the idea that phenotypic outcomes for polygenic traits can be reached through different combinations genetic variants due to genetic redundancies ^99^.

**Figure 6.**
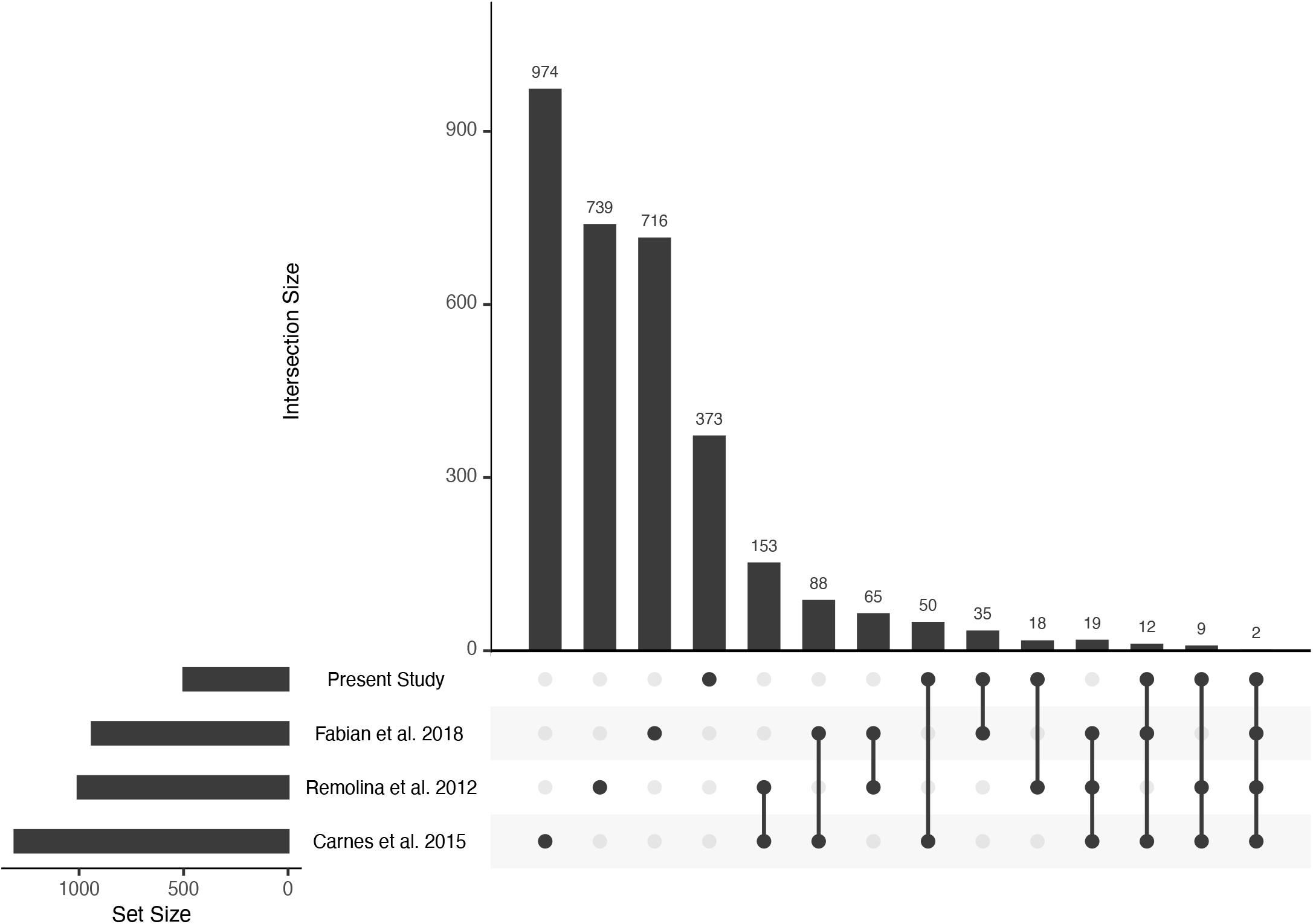
An UpSet plot showing patterns of overlap between candidate genes from the present study, Remolina et al. 2012, Carnes et al. 2015, and Fabian et al. 2018.

### Conclusions

Broadly speaking, our effort to incorporate metabolomics into the E&R framework has successfully generated insights into the factors underlying longevity differences in this experimental system. On its own, comparing and contrasting metabolomic profiles between our experimental populations revealed a number of key mechanisms shaping aging and longevity differences across the system. Our findings also suggest that, with regards to longevity, the relationship between the genome and the metabolome is complex and highly polygenic. However, the incorporating metabolomic results into our genomic analysis allowed us to narrow our focus to a tractable, and potentially more impactful, subset of candidate genes. The presence of key established aging genes and enrichment for functional gene clusters and networks related to longevity also suggests we are capturing some meaningful aspect of the genetic mechanisms underlying longevity differences in this system. Taken together, this study serves as a proof of concept that combining different types of omics data in the E&R context may have significant benefits when attempting to parse the physiological and genetic mechanism shaping complex phenotypes like longevity.

Our work also adds to the growing body of evidence that canonical genetic mechanisms are not always the primary drivers of complex trait variation in “real” populations or laboratory population approximating real populations ^28,29,120^. While these mechanisms are highly relevant in certain contexts (i.e. mutant screens, studies in specific genetic backgrounds, etc.), reality is more complex in outbred populations and other sources of variation with perhaps smaller individual effect sizes are more relevant. And similar functional outcomes can be achieved through different genetic mechanism. As such, there is a clear need for approaches specifically designed to contend with this complexity when seeking to understand complex trait variation in real populations.

## Acknowledgments

We would like to thank our undergraduate colleague Kit Neikirk and Taylor A Rodman for helping to optimize the analysis technique. We would also like to thank Dr. Michael R. Rose for access to the experimental populations maintained in his lab at UC Irvine. This work was supported by The Burroughs Wellcome Fund Award Career Awards at the Scientific Interface (CASI), UNCF/BMS EE Just Faculty Fund, the Ford Foundation, and NIH SRP subaward to #5R25HL106365-12 from the NIH PRIDE Program all awarded to AJH. MRM. is supported by the Burroughs Wellcome Fund and Howard Hughes Medical Institute via the PDEP and Hanna H. Gray Fellows Program. Dr. Melanie McReynolds, Dr. Edgar Garza Lopez, and Dr. Derrick J. Morton all provided consulting for this project.

## Data Availability

Core data files (metabolite readings, SNP tables, results of statistical analyses, etc.) are available through Dryad (https://doi.org/10.5061/dryad.547d7wm92), and scripts used to carry out analysis linking genomic results to metabolomic results are available through Github (https://github.com/ttbarter317/Fruit-fly-Genomics-and-Metabolomics).

## Institutional Review Board Statement

Not applicable.

## Informed Consent Statement

Not applicable.

## Author Contributions

Conceptualization MAP and AJH

Methodology: KRA, MAP, ZV, AJH

Validation: MAP, ZV, HKB, AGM, MRM

Formal Analysis: MAP, TTB, HKB, KRA

Investigation: MAP, ZV, HKB, AGM, MRM, AJH,

Resources: MAP, AJH

Writing—Original Draft Preparation, MAP, AGM, HKB, ZV, MR, AJH

Writing—Review & Editing MAP, KRA, ZV, HKB, AGM, DJM, MRM, AJH

Visualization, MAP, ZV, HKB, AGM

Supervision, MAP, AJH

Project Administration MAP, AJH

Funding Acquisition AJH All authors have read and agreed to the published version of the manuscript.

## Supplementary Figures

**Supplementary Figure 1.**
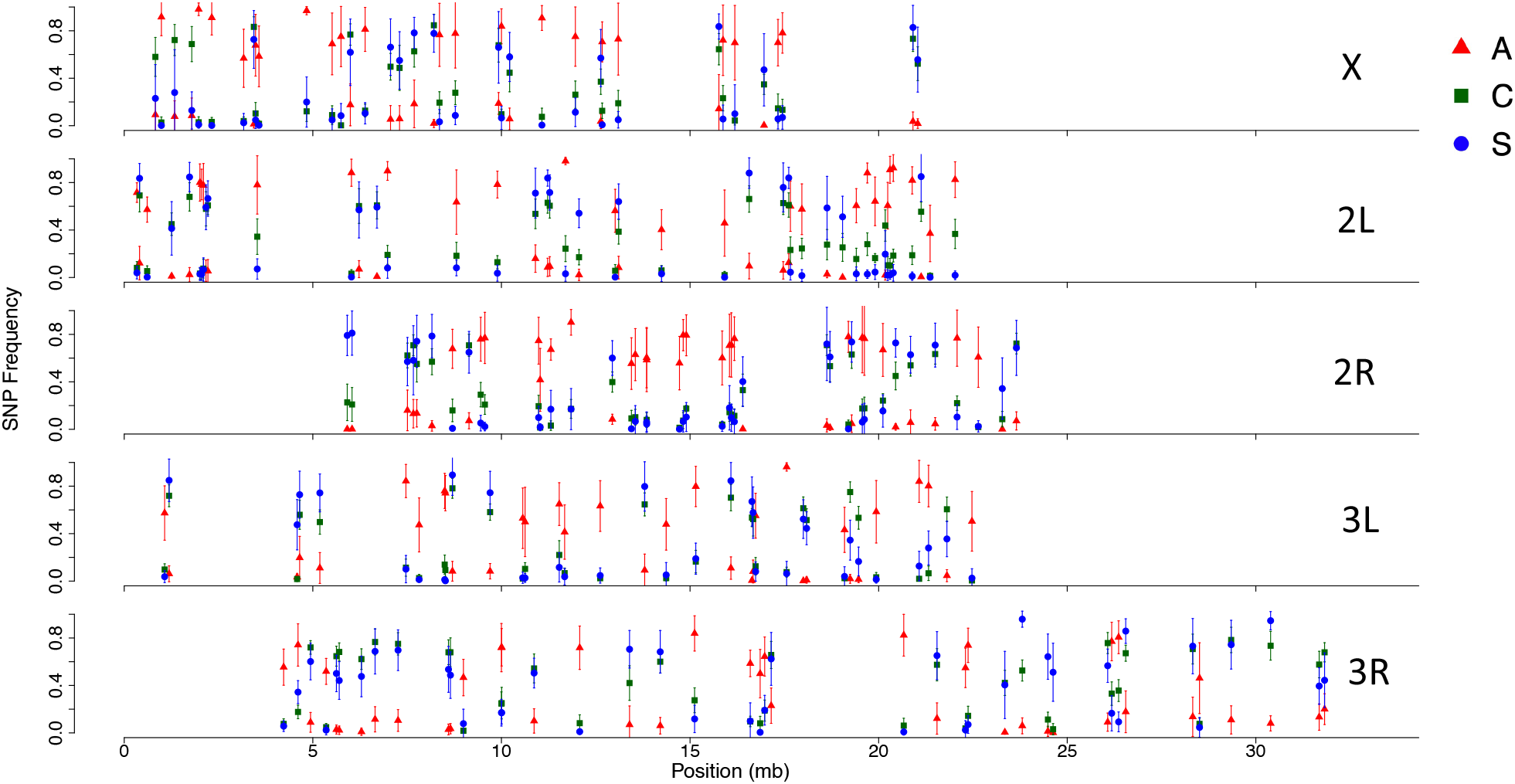
Mean SNP frequencies in the A, C, and S populations at sites that best predict patterns of candidate metabolites differentiation based on FLAM analysis. Panels represent the major chromosome arms and error bars are based on standard deviation.

**Supplementary Figure 2.**
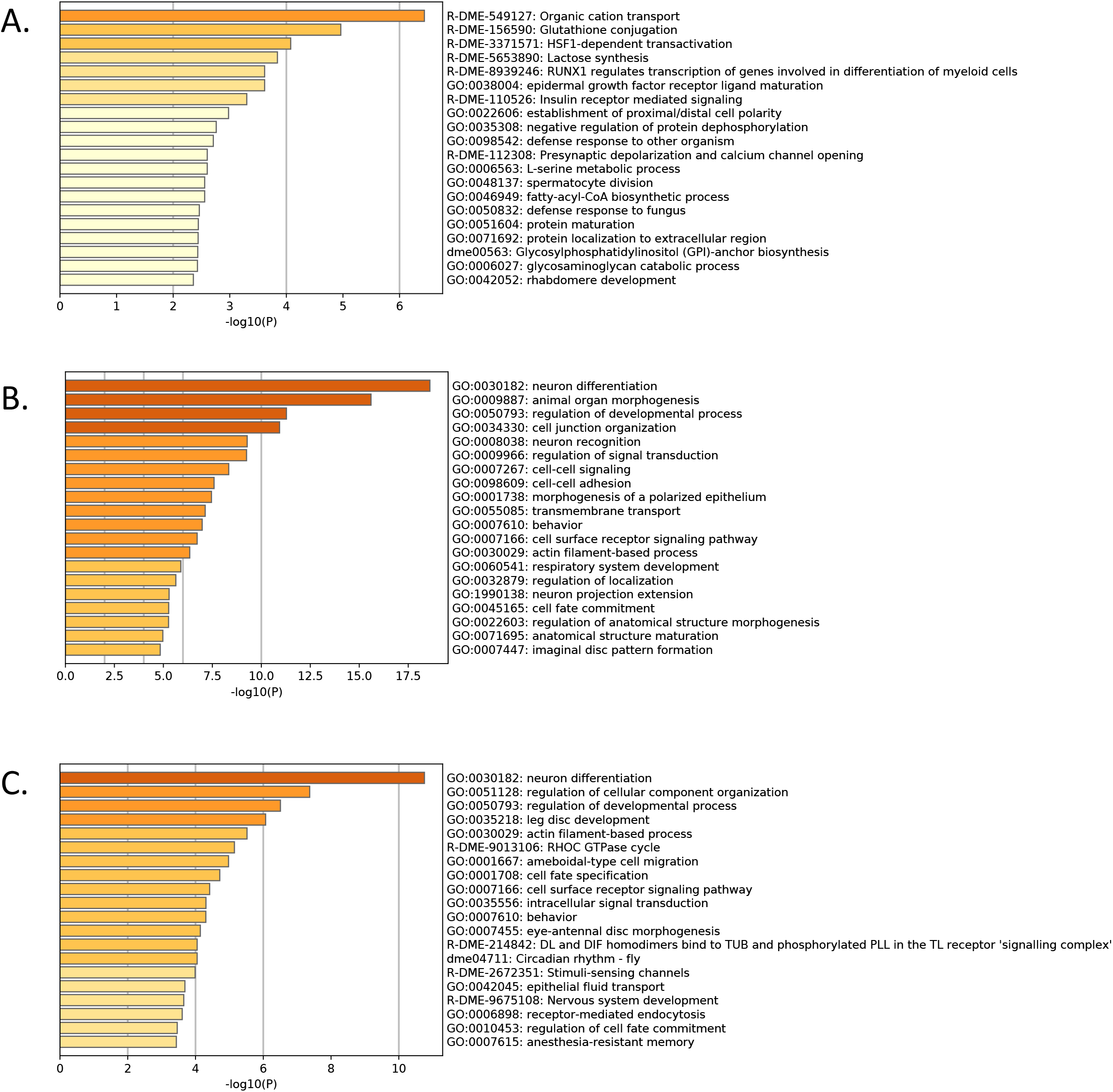
Heatmap of top 20 enriched GO clusters based on candidate longevity genes identified by Remolina et al. 2012 (A), Carnes et al. 2015 (B), and Fabian et al. 2018 (C).

